# MNase stratification reveals heterogeneous 5hmC in naive B cells

**DOI:** 10.1101/2025.08.24.672045

**Authors:** Michael Hsu, Heng-Yi Chen, Fang-Yun Lay, Qin Ma, Ben Delatte, Chan-Wang Jerry Lio

## Abstract

DNA demethylation is essential for gene activation and is primarily mediated by the Ten-Eleven-Translocation (TET) dioxygenase family. TET initiates the demethylation by oxidizing 5-methylcytosine (5mC) to 5-hydroxymethylcytosine (5hmC), a chemically stable derivative that is not only an intermediate in demethylation but also an epigenetic mark. 5hmC is enriched at active gene bodies, promoters, and enhancers that exist at accessible chromatin. Yet, 75-90% of genomic DNA is physically packaged in nucleosomes, raising the question about where 5hmC resides relative to nucleosomes. To address this question, we used micrococcal nuclease (MNase) to biochemically stratify 5hmC according to nucleosome protection in naive mouse B cells. Using multiple approaches, we showed that ∼95% of 5hmC peaks are retained after MNase digest, suggesting that 5hmC is nucleosome-associated. Despite the majority of 5hmC being similar in nucleosomal DNA compared to total DNA, we identified minor subsets that were MNase-sensitive (accessible 5hmC, ∼3%) or further MNase-resistant (protected 5hmC, ∼1-2%). Integrative analyses revealed that the accessible 5hmC is preferentially located at promoters and enhancers, and the presence of promoter-proximal accessible 5hmC showed a stronger correlation with gene expression than total 5hmC. In contrast, protected 5hmC is enriched in chromatin states with mixed active and repressive features. These results shed new light on the relationship between 5hmC and accessible chromatin, suggesting 5hmC is predominantly associated with nucleosomes and revealing heterogeneous 5hmC subsets with potential distinct regulatory roles.

## Introduction

DNA methylation, primarily at the CpG dinucleotides, is a fundamental epigenetic modification that regulates gene expression, genome stability, and cell fate decisions. In mammals, unmethylated cytosines are methylated by the *de novo* DNA methyltransferases DNMT3A and DNMT3B into 5-methylcytosine (5mC).

DNMT1 maintains the methylation pattern during replication^1,2^. In contrast, DNA demethylation is mediated by the TET (Ten-Eleven Translocation) dioxygenase family (TET1, TET2, and TET3), which sequentially oxidizes 5mC to 5-hydroxymethylcytosines (5hmC). Once thought to be merely a transient intermediate, 5hmC is now recognized as a stable and functionally distinct epigenetic mark enriched at active gene bodies, enhancers, and promoters^3–6^. TET enzymes are essential for cell differentiation and the functions of immune cells, including B lymphocytes, as we and others previously described^7–11^. Furthermore, *TET2* is frequently mutated in hematological malignancies such as diffuse large B cell lymphomas and myeloid leukemias, highlighting their role as essential tumor suppressors^12^.

In mammalian cells, 75-90% of the genomic DNA is packaged into nucleosomes, which are protein complexes of approximately 146 base pairs of DNA and histone octamers^13,14^. Histone post-translational modifications provide another layer of epigenetic regulation critical for gene regulation and chromatin structure. DNA methylation is generally associated with nucleosomal DNA and repressive chromatin, whereas 5hmC has been more strongly linked with accessible chromatin and transcriptional activity. A previous study suggested that TET1 and 5hmC are associated with the labile, MNase-sensitive nucleosomes^15^. However, the extent to which 5hmC is nucleosome-bound and how such localization influences chromatin states remained unclear.

In this study, we set out to define the chromatin context of 5hmC in naïve mouse B cells. By integrating micrococcal nuclease (MNase) to digest the accessible DNA and obtain the nucleosomal DNA coupled with genome-wide 5hmC profiling, we distinguished protected (MNase-resistant) versus accessible (MNase-sensitive) 5hmC and compared their enrichment across chromatin states. We hypothesize that while most 5hmC is accessible, some will associate with nucleosomes and may have diverse functions in gene regulation. Unexpectedly, we found that the vast majority of 5hmC resides in MNase-resistant DNA, suggesting potential nucleosome association. The protected 5hmC closely mirrors the genomic distribution of total 5hmC. Nonetheless, a minority of regions displayed differential MNase sensitivity, corresponding to functionally distinct 5hmC subpopulations. The more accessible 5hmC showed strong enrichment at active promoters and enhancers, whereas protected 5hmC was preferentially associated with bivalent-like states marked by both H3K9ac and H3K9me3. Remarkably, the promoter-proximal accessible 5hmC showed a stronger correlation with gene expression than the protected 5hmC, suggesting that nucleosome context diversifies the regulatory potential of 5hmC. Together, our findings challenge the prevailing view that 5hmC primarily marks accessible DNA and instead reveal that most 5hmC is potentially nucleosome-bound, existing in distinct chromatin environments that may differentially shape transcriptional regulation.

## Results

### Most 5hmC resides within nucleosome-protected DNA

5hmC is known to be enriched at active gene bodies, promoters, and enhancers. All these regions often exist in accessible and transcriptionally permissive chromatin^16^. Therefore, the prevailing expectation is that most 5hmC should be enriched in accessible DNA. Yet, since most eukaryotic DNA (∼75%-90%) is wrapped in nucleosomes^13,14^, we hypothesized that a fraction of 5hmC is on nucleosome-bound DNA that may have an alternative function compared to the 5hmC at open chromatin. To directly address this issue, we used murine naïve B cells as the model and biochemically stratified 5hmC by chromatin state using micrococcal nuclease (MNase). MNase can preferentially digest accessible DNA without significantly affecting most nucleosome-bound DNA. Importantly, MNase provides a physical partition of 5hmC by nucleosome association that is complementary to accessibility assays such as ATAC-seq (which report on exposed DNA rather than protected DNA). To ensure obtaining mononucleosomes, we titrated MNase digestions and selected the highest concentration that was able to remove both accessible DNA and labile nucleosomes (**Fig. S1**). As the control, we sonicated purified DNA to the size of nucleosomal DNA (∼147bp). This fraction should encapsulate all 5hmC regardless of chromatin states. The 5hmC enrichment in each set of the DNA samples was analyzed using a modified hMe-Seal-seq, a pulldown method mediated by T4 β-glucosyltransferase and CLICK chemistry^17^. Herein, we refer to the 5hmC captured from MNase-derived mononucleosomes as “**nucleosomal 5hmC**” and from the size-matched sonicated control as “**total 5hmC**.”

If most 5hmC is localized in the MNase-sensitive accessible DNA, we expected that MNase treatment would remove a significant portion of 5hmC and reveal the locations of 5hmC that are nucleosome-bound. To our surprise, we found that the 5hmC distributions in the genome are highly similar between total and nucleosomal 5hmC, with 95.4% of the peaks shared between the two fractions (**Fig. 1B–C; common 5hmC**). We noticed a minority were selectively **protected** (1.6%; higher MNase resistance) or **accessible** (3%; lower MNase resistance; **Fig. 1B-C**), although they primarily differed only by levels of enrichment rather than their absence or presence (see below). As most 5hmC peaks are not sensitive to MNase, this result suggests that most 5hmC is not accessible, challenging our original presumption that most 5hmC is accessible.

**Figure 1.**
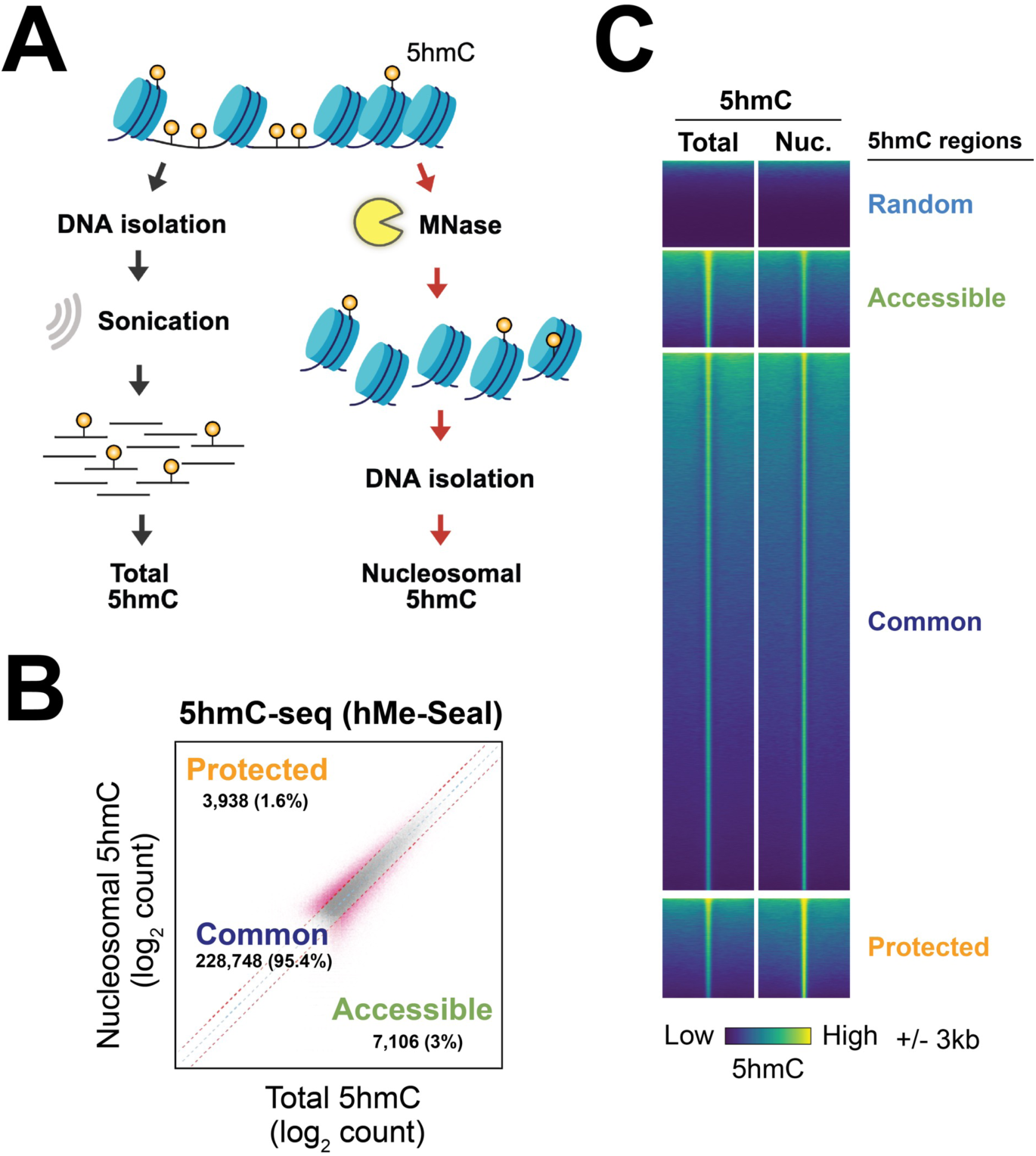
The majority of 5hmC is located within MNase-resistant DNA. **(A)** Schematic for analysis of 5hmC in total and Nucleosomal DNA. To analyze the total 5hmC, genomic DNA from naïve mouse B cells was isolated and sonicated to ∼150bp (similar to the size of nucleosomal DNA). To obtain nucleosomal DNA, nuclei from naïve mouse B cells were digested using micrococcal nuclease (MNase), followed by DNA purification. The genome-wide distribution of 5hmC in total and nucleosomal DNA was analyzed using hMe-Seal-seq (see Methods). **(B)** 5hmC enrichment patterns are similar between total and nucleosomal DNA. A scatterplot of log_2_ 5hmC counts in Total (x-axis) and Nucleosomal (y-axis) 5hmC samples is shown. Differential 5hmC regions were defined with a 1.5-fold difference (fuchsia lines) and FDR ≤ 0.05. Red dots represent the differential 5hmC regions: regions preferentially enriched in total 5hmC were considered “Accessible,” and those enriched in nucleosomal 5hmC were defined as “Protected.” The shared regions that did not meet the statistical cut-off were defined as “Common.” **(C)** 5hmC enrichment at the differential regions. Representative heatmaps of total and nucleosomal 5hmC enrichment over peaks of accessible, common, or protected 5hmC are shown. Randomized regions were demonstrated as the controls to demonstrate specific enrichment of 5hmC. All regions are centered and display enrichment +/- 3kb from the centers. Three biological replicates were analyzed for each DNA preparation, and the average signals are shown in (B) and (C).

To test whether MNase digestion causes a global loss of 5hmC signal, we added a fixed amount of 5hmC-rich DNA from T4gt phage as an exogenous spike-in control before library preparation and sequencing. The phage genomes are heavily modified with 5hmC and are routinely used as positive controls for 5hmC pulldown, as in hMe-Seal–style assays. The captured DNA was sequenced, and the spike-in read counts were normalized to total depth. Under an accessibility-dominant model, MNase digestion would deplete endogenous (MNase-sensitive) 5hmC, thereby increasing the spike-in fraction (**Model 1 in Fig. 2C**); under a nucleosome-dominant model, MNase would have a limited effect on endogenous 5hmC, and the spike-in fraction would remain similar to the ratio of spike-in to total 5hmC (**Fig. 2B**; **Model 2 in Fig. 2C**). Using an exogenous reference provides quantitative sensitivity to global shifts that standard depth normalization can mask, analogous to ChIP-Rx spike-in normalization for chromatin profiling^18^. Our results showed that the normalized T4gt read counts were similar between those mixed into the total 5hmC and nucleosomal 5hmC (**Fig. 2D**). Thus, the result was more consistent with the nucleosome-dominant model (model 2). Finally, to further validate the above results, we use mass spectrometry to directly quantify the abundance of 5hmC in the sonicated and MNase-treated DNA. The results showed that both sonicated and MNase-digested DNA have a similar amount of 5hmC (**Fig. 2E**). Therefore, different from the prevailing assumption that 5hmC is primarily associated with accessible DNA, our data suggest that most 5hmC is potentially nucleosome-bound.

**Figure 2.**
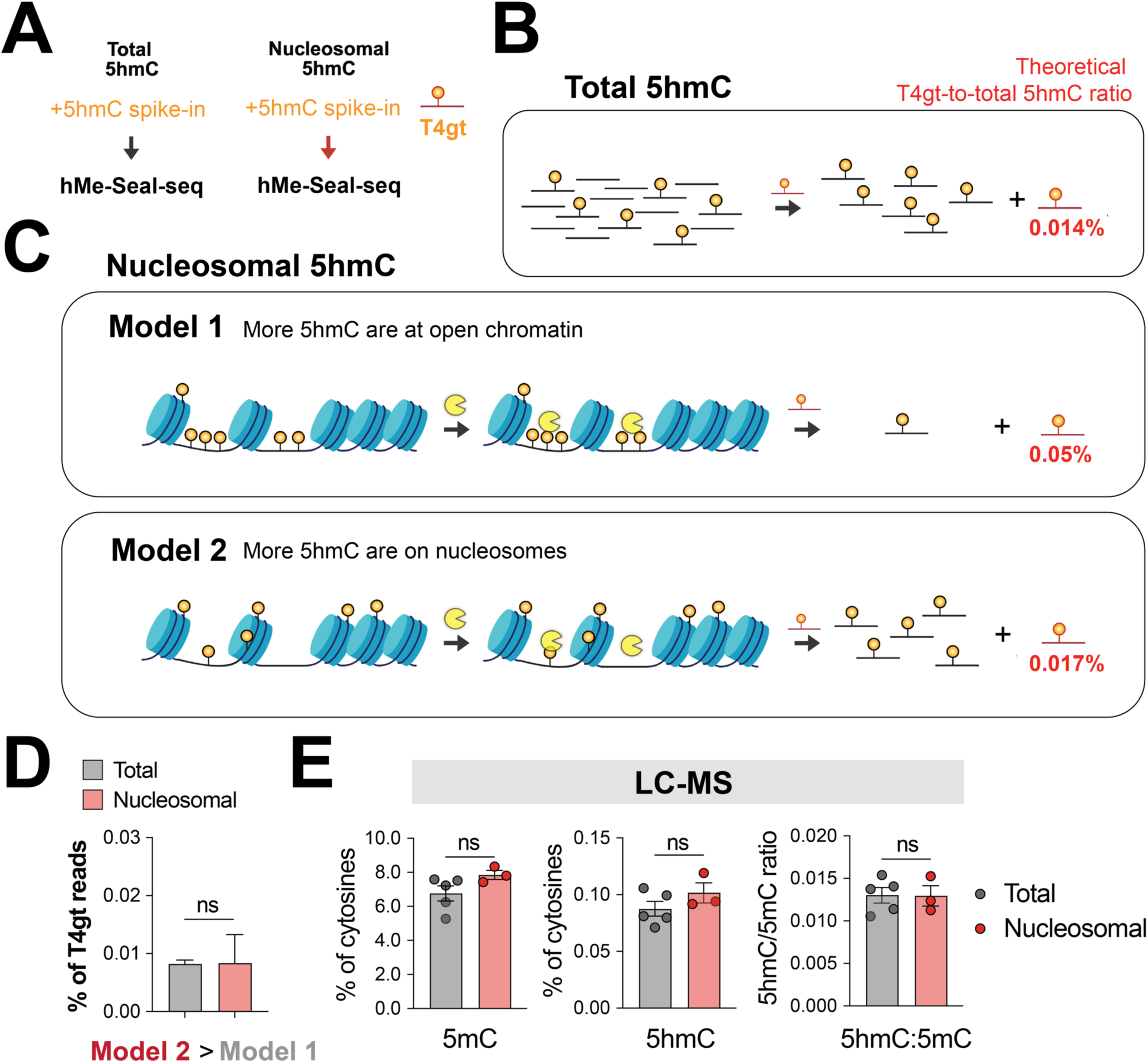
5hmC levels are similar between total and nucleosomal DNA. **(A)** Schematic for the inclusion of 5hmC spike-in. T4gt phage DNA from that containing 100% 5hmC at all cytosine positions was added to all samples before 5hmC enrichment analysis using hMe-Seal-seq. **(B-C)** Diagrams of two alternative 5hmC distribution models. (B) 5hmC-containing DNA in total DNA and spike-in was enriched, and the theoretical T4gt-to-total 5hmC ratio is shown. (C) Two models of 5hmC distributions and predictions. In Model 1, if more 5hmC exists at open chromatin, most of these 5hmC will be removed by MNase, resulting in a higher T4gt-to-total 5hmC ratio relative to that in (B). In Model 2, if the majority of 5hmC is on nucleosomes, most of the 5hmC will remain even after MNase digestion. As a result, the T4gt-to-total 5hmC ratio will be similar to (B). **(D)** T4gt- to-total ratios are similar regardless of MNase digestion. Mapped T4gt reads were normalized to total reads to obtain the %T4gt reads as described in Methods. Note: the results are more consistent with the prediction from Model 2, suggesting 5hmC is more likely to be on nucleosomal DNA. **(E)** Confirmation of 5hmC levels using mass spectrometry. The 5hmC, 5mC, and C (cytosines) on total genomic and nucleosomal DNA were analyzed using liquid chromatography - mass spectrometry (LC-MS). The percentage of 5mC, 5hmC, and the ratios of 5hmC among 5mC were shown. Two independent experiments and a representative experiment is shown.

### Accessible 5hmC is linked to higher chromatin accessibility and lower DNA methylation

While the majority of the 5hmC is MNase-resistant, we noticed a total of around 5% regions that are relatively more MNase-sensitive (accessible 5hmC) and more resistant (protected 5hmC) (**Fig. 1B**). To determine if MNase sensitivity can be used to stratify subpopulations of 5hmC-enriched areas that may exist at different chromatin environments, we sought out to correlate the differential 5hmC enrichment with chromatin accessibility (**Fig. 3**). Consistent with the previous observations, the enrichment of 5hmC is generally associated with increased chromatin accessibility (**Fig. 3A-B**). The average chromatin accessibility is directly correlated with the MNase sensitivity, where the accessible 5hmC shows higher accessibility, whereas protected 5hmC shows lower accessibility (**Fig. 3A-B**). The regions flanking the 5hmC peaks are more accessible relative to the centers, consistent with the notion that 5hmC is potentially nucleosome-bound (**Fig. 3B**).

**Figure 3.**
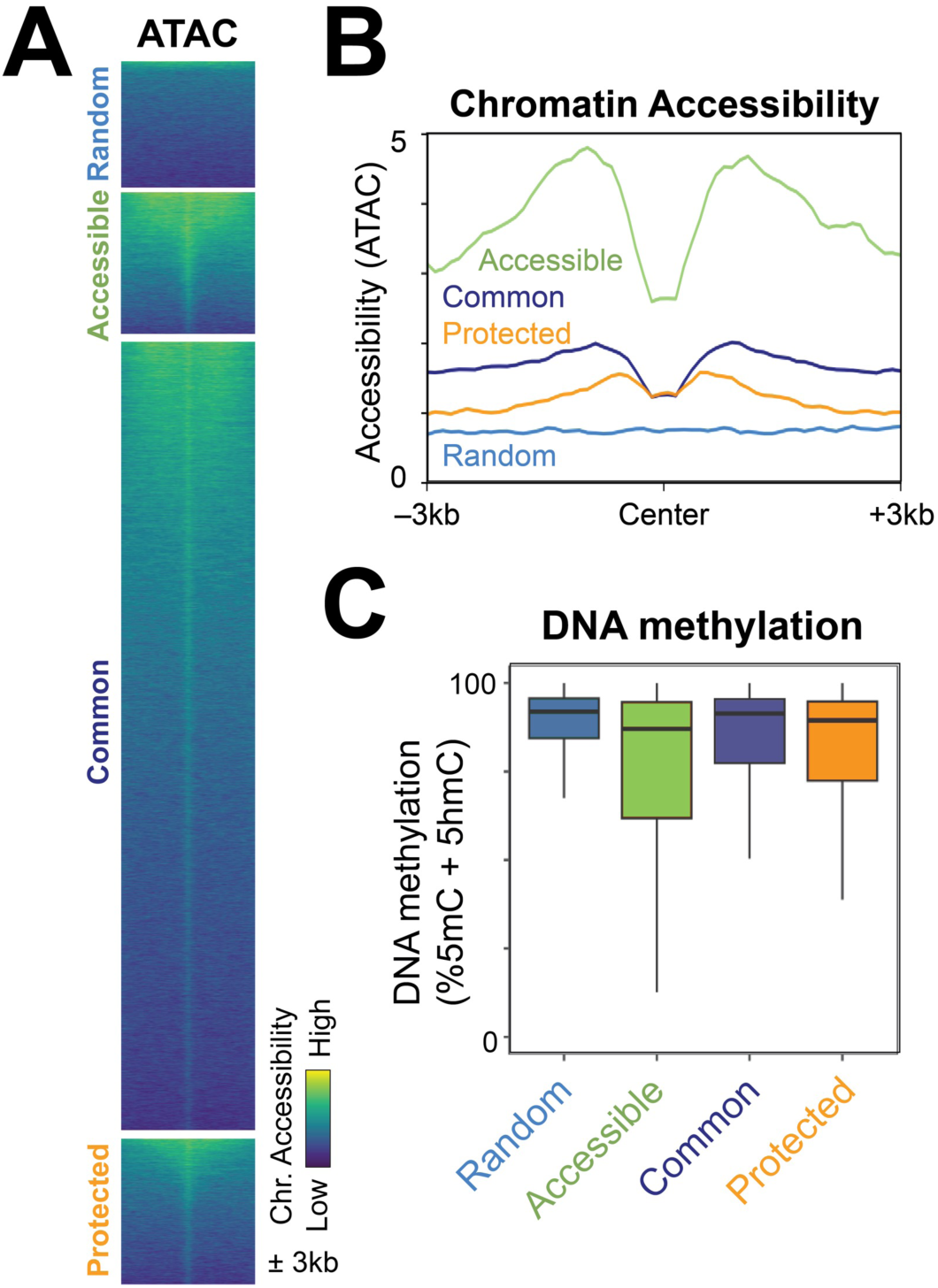
Accessible 5hmC is linked to higher chromatin accessibility and lower DNA methylation. (A-B) Chromatin accessibility around differential and common 5hmC regions. Chromatin accessibility in naïve B cells was analyzed from our previously published ATAC-seq data. The accessibility in the indicated regions (+/- 3kb) is shown as heatmaps (A) and profile plots (B). (C) Accessible 5hmC regions have decreased DNA methylation. CpG methylation levels obtained from genome-wide bisulfite sequencing at +/-500bp of the 5hmC peaks are shown as box plots. Note that bisulfite sequencing cannot distinguish 5mC versus 5hmC.

We next asked whether these 5hmC subsets occupy distinct methylation environments. Using the data from a matched whole-genome bisulfite sequencing, we quantified CpG methylation across ±500 bp windows surrounding 5hmC peak centers. Consistent with prior studies, we show that chromatin accessibility is anticorrelated with DNA methylation (**Fig. 3B-C**). The accessible 5hmC peaks displayed markedly reduced methylation (**Fig. 2C**), whereas the protected 5hmC peaks retained relatively higher methylation levels (**Fig. 2C**). Together, the analyses indicate that 5hmC is not at a uniform epigenetic state: accessible 5hmC marks hypomethylated regions, while the protected 5hmC may persist in more methylated, alternative chromatin environments.

### Distinct transcription factor (TF) contexts for accessible and protected 5hmC

To explore if the 5hmC subsets may be regulated by different regulatory networks, we analyzed the TF motifs that are enriched. Consistent with our previous finding, 5hmC-containing regions are enriched for selected transcription factors, particularly the ETS family members such as PU.1, which has been shown to recruit TET enzymes (**Fig. S2**). To identify the transcription factors associated with the accessible and protected 5hmC, we used the motifs in the common 5hmC regions as background. The results showed an interesting distinction in TF families. The accessible 5hmC regions are preferentially enriched in ETS (PU.1) and IRF, consistent with the observation that PU.1 and E2A/IRF cooperate and recruit TET proteins to enhancers during B cell differentiation^8,19^. In contrast, the protected regions are enriched in the KLF family proteins (**Fig. 4**). This observation is consistent with the known function of some KLF proteins, including KLF4 and KLF1, which can function as pioneer transcription factors that can bind to nucleosome-bound DNA^20,21^. Interestingly, we noticed that several motifs contain CpG sites (**Fig. 4A, 4C**). CpG methylation is known to modulate TF–DNA affinity in a context-dependent manner. For instance, KLF4 can recognize or even prefer methylated CpG in its binding site, whereas numerous other TFs, including other KLFs and IRFs, are inhibited by CpG methylation^22,23^.

**Figure 4.**
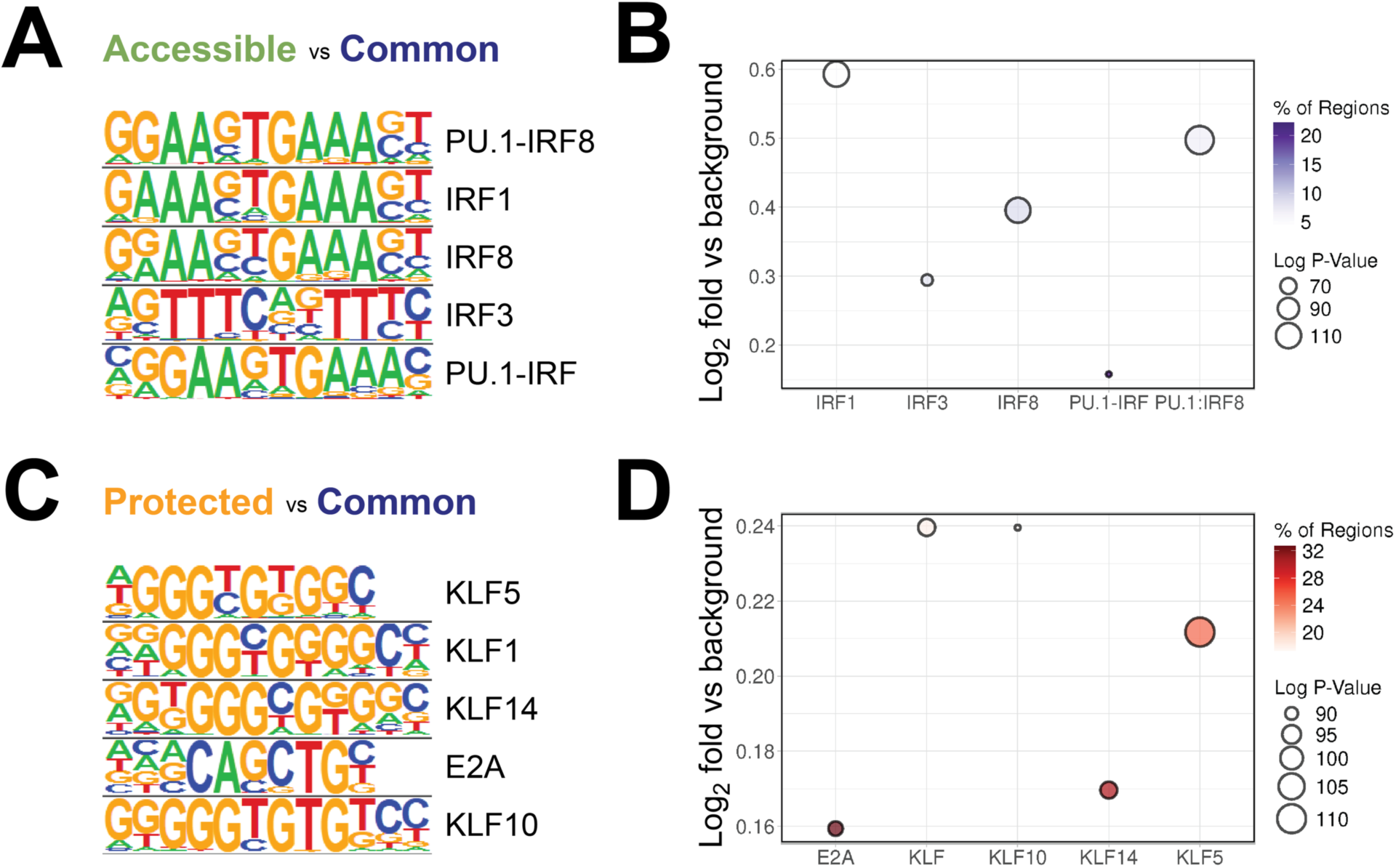
Distinct transcription factor (TF) contexts for accessible and protected 5hmC. (A-D) Motif enrichments at accessible (A-B) and protected (C-D) 5hmC regions were analyzed using the common 5hmC regions as the background. The sequences (A, C) and enrichment levels (B, D) for top five motifs for each region subset are shown.

Together, these data support a model in which accessible 5hmC coincides with the PU.1/IRF regulatory network, while the protected 5hmC is preferentially associated with the KLF family. Whether 5hmC deposition promotes TF occupancy in these settings or is downstream of TF binding remains an open question.

### Accessible 5hmC at promoters strongly correlates with transcriptional activity

To understand if the differential 5hmC regions are preferentially enriched at distal or proximal locations, We annotated each 5hmC peak as proximal or distal based on its distance to the nearest transcription start site (TSS; **Fig. 5A**). In both MNase-sensitive (“accessible 5hmC”) and MNase-resistant (“protected 5hmC”) subsets, the vast majority of peaks were distal (>95%), with ∼4–5% proximal, slightly exceeding the proximal fraction in the common 5hmC set (∼3.2%). To test whether these subsets track gene expression, we ranked genes by the expression levels and partitioned them into quartiles (Q1–Q4) and linked each 5hmC peak to the nearest gene. As expected from prior studies, promoter-proximal 5hmC associated positively with transcription compared to promoter-matched randomized controls (**Fig. 5B, left**)^24^. Strikingly, promoters with accessible 5hmC showed significantly higher expression than those with common 5hmC, whereas promoters with protected 5hmC expressed at levels indistinguishable from random, indicating little or no positive association (**Fig. 5B, left**). In contrast, all distal 5hmC did not improve expression prediction relative to random regions (**Fig. 5B, right**). The results suggest that 5hmC at different chromatin contexts and locations can have varying correlations with transcriptional activity. Promoter-proximal accessible 5hmC correlates with higher gene expression, whereas promoter-proximal protected 5hmC did not show a robust association with expression and was comparable to promoter-matched random controls, suggesting limited predictive value in this setting.

**Figure 5.**
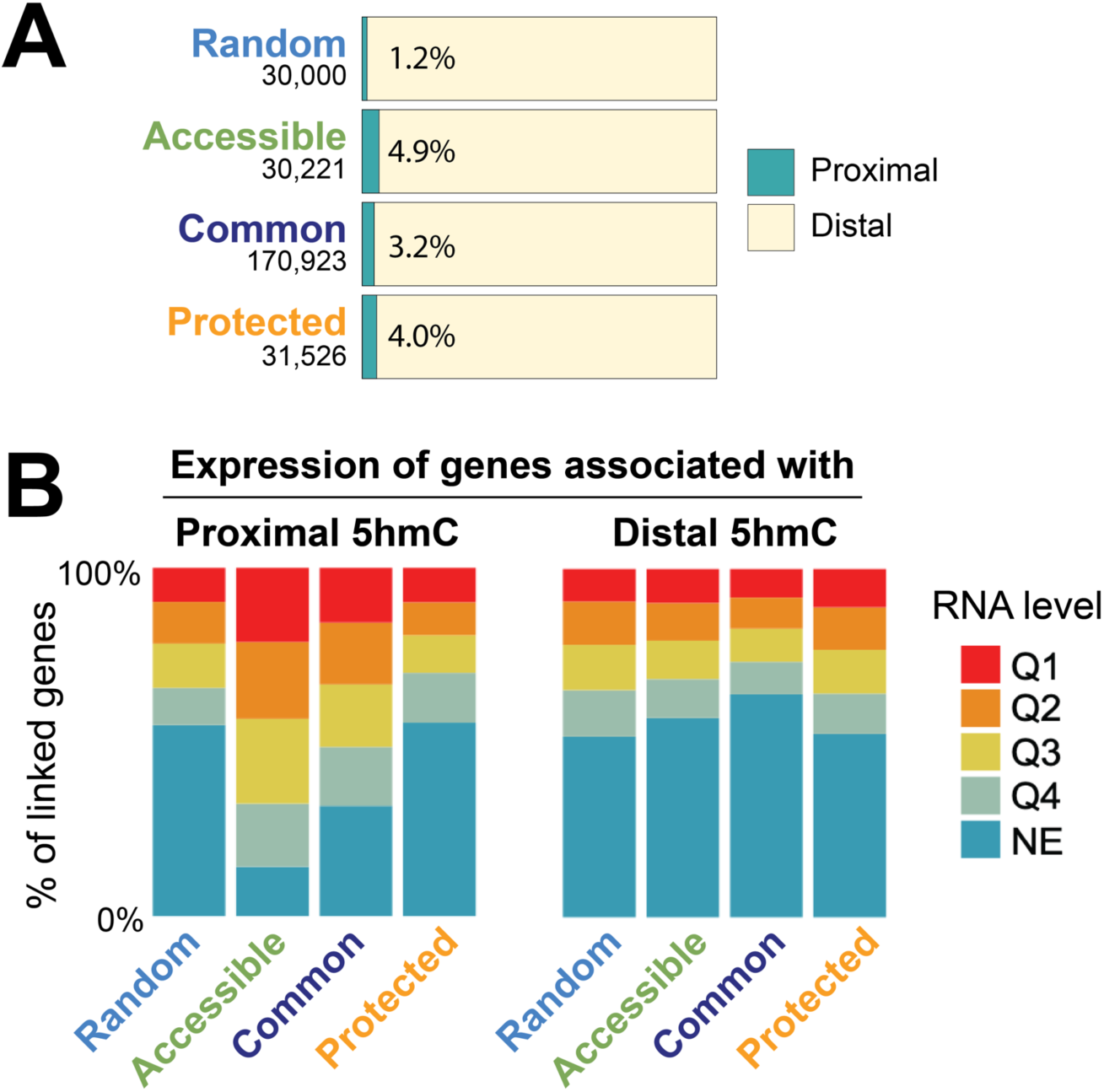
Accessible 5hmC at promoters strongly correlates with transcriptional activity. **(A)** Distribution of regions relative to promoters. Regions were considered to be proximal if they were within +/- 500bp relative to gene promoters. **(B)** Gene expression in naïve B cells was first categorized into five groups, with Q1 being the 25% highest expressed and Q4 the lowest expressed genes. NE indicated genes that were not expressed. Each region within each 5hmC subset or random regions was assigned to the closest genes. The expression levels for all the genes linked to each subset were tabulated.

### Chromatin-state stratification reveals heterogeneity among 5hmC subsets

To systematically characterize these subsets, we trained a 20-state ChromHMM model on naïve B-cell ChIP-seq for histone marks and chromatin regulators (45 datasets), then annotated states by comparing their emission profiles to reference “universal” mouse annotations (**Fig. 6A and S3**)^25–27^. Fig. 6A shows the probability of each epigenetic mark to be found in each state (row), and each category is labeled by colors (“Chr. States”; see legend in **Fig. 6C**). For instance, most of the typical promoter states have varying degrees of H3K4me3 and H3K27Ac (state 1-4; colored red). The analysis showed that many of the bins in the genome are annotated as “other” states, which include quiescent and no-enrichment regions (“Genome”; heatmap in **Fig. 6B** and same data as bar graph in **6C**). The analysis for random regions showed a similar result, where most of them overlapped with the “others” states (“Random”, **Fig. 6B-C**). As a control, we analyzed the chromatin states for all the accessible regions identified via ATAC-seq in naïve B cells. Most of the accessible regions (peaks) overlapped with promoter states, especially states 1 and 2 (“ATAC peaks”, **Fig. 6B-C**), consistent with the known relationship between open chromatin and regulatory DNA.

**Figure 6.**
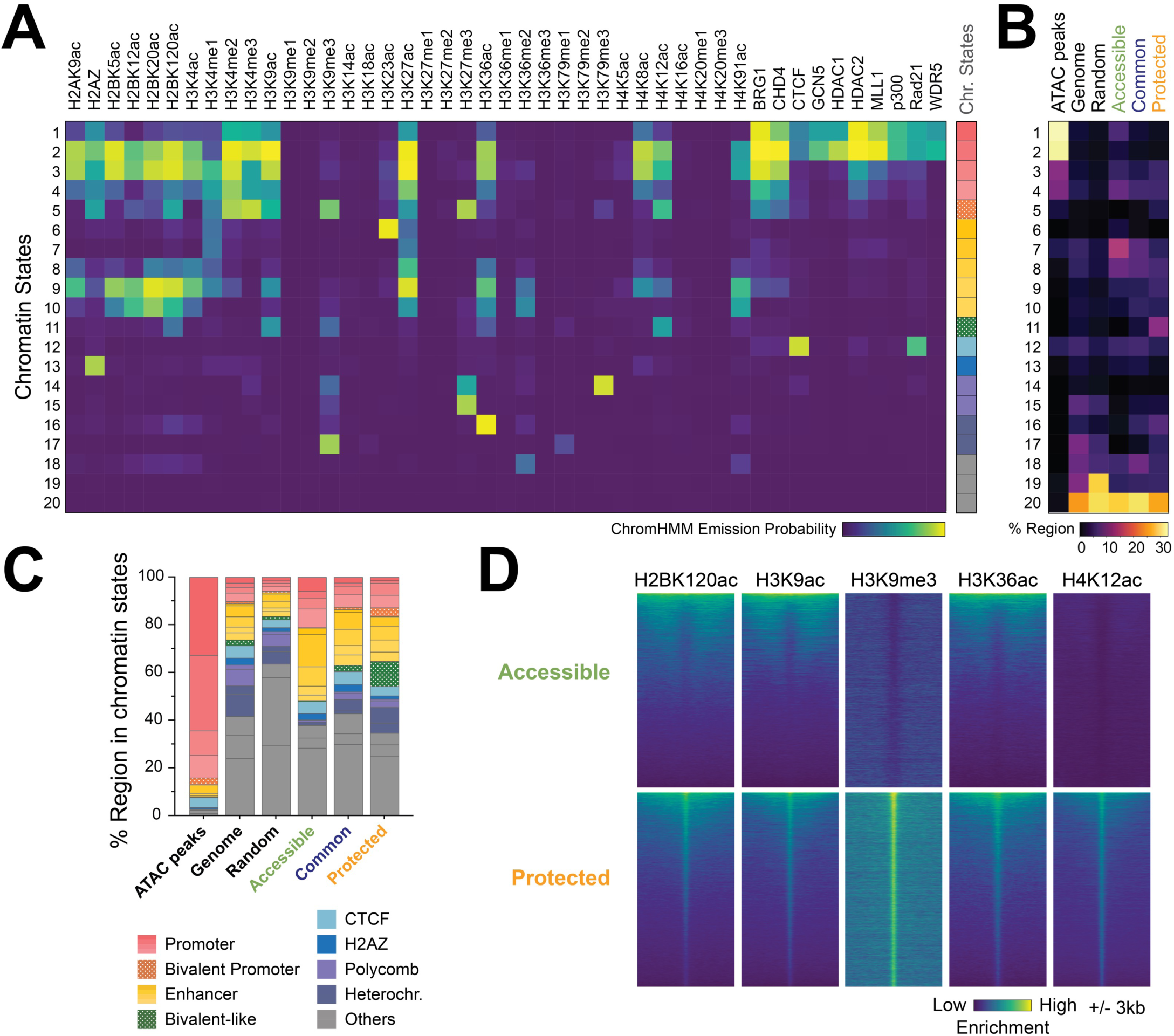
Chromatin-state stratification reveals heterogeneity among 5hmC subsets. **(A)** Chromatin state assignment. 45 published ChIP-seq datasets from mouse naïve B cells were used to train ChromHMM to generate 20 unique chromatin states. The genome was divided into 200bp bins, and each was annotated for a chromatin state. The emission probabilities (chance of appearing in a specific state) for each chromatin state (row) are shown. The identity of each state was annotated (**Fig. S**) and labeled with colors on the right panel (see legend in **C**). **(B-C)** Distributions of 20 chromatin states among control, genomic, 5hmC subsets, and accessible regions are shown as a heatmap (B) and stacked bars (C). **(D)** Selected histone marks related to bivalent-like states. Heatmaps showing the enrichment of 5 histone marks at accessible and protected regions. Heterochr., heterochromatin.

**Figure 7.**
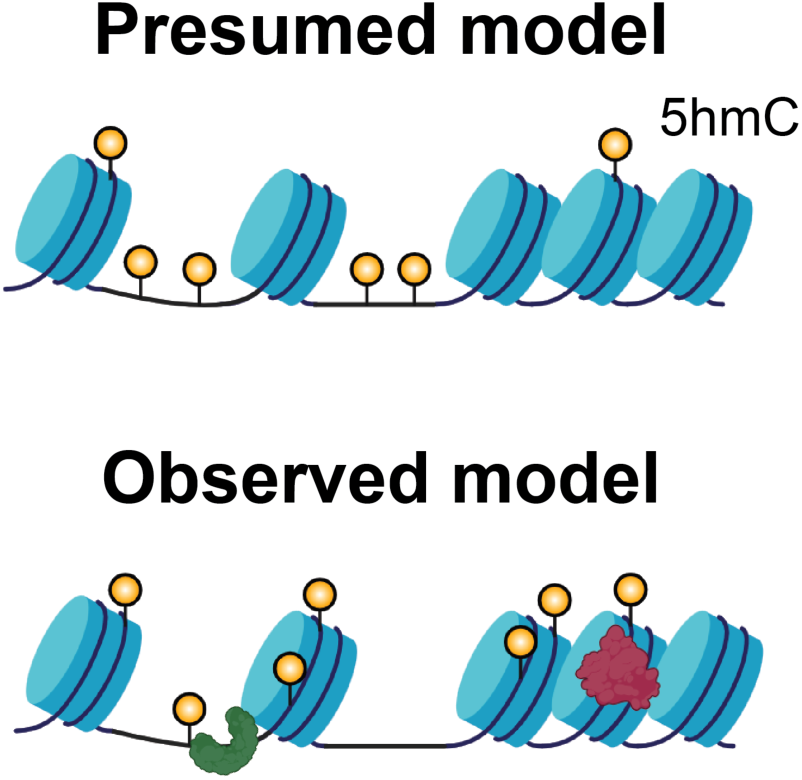
Current working model. While 5hmC was previously shown to be more abundant in accessible regions (presumed model), the data here suggest that 5hmC is more likely to be nucleosome-bound. Moreover, 5hmC-enriched regions are not homogeneous, and they may function differently according to the local chromatin context and potentially the involvement of different TF families.

Next, we ask if the differential 5hmC regions preferentially associate with specific chromatin states. We found that all three classes show significant enrichment of promoter (red) and enhancer states (yellow; **Fig. 6B-C**), with the accessible 5hmC showing increased overlap (**Fig. 6B-C**). By contrast, the protected 5hmC has significantly higher overlaps with mixed active and repressive features (“bivalent-like”) and in H3K9me3-rich heterochromatin-associated states (**Fig. 6C-D**). We note that “bivalency” is usually defined by the presence of H3K4me3 and H3K27me3 at the same locus. The “bivalent-like” state here means the concurrence of the active H3K9ac and repressive H3K9me3 (**Fig. 6**). Consistent with increased H3K9me3, the protected 5hmC also has more overlap with heterochromatin states compared to the accessible 5hmC (**Fig. 6D**).

Altogether, our data demonstrated that most 5hmC is potentially nucleosome-bound. Nonetheless, heterogeneity exists among 5hmC-enriched regions, which can be stratified by their degree of association with nucleosomes. The more accessible 5hmC (MNase-sensitive) at the promoters is a better indicator for gene expression. The more protected 5hmC (MNase-resistant) is preferentially embedded in bivalent-like/heterochromatin-associated states, consistent with poised or repressive chromatin contexts.

## Discussion

In this study, we set out to investigate whether 5hmC exists on nucleosomes. Surprisingly, contrary to the prevailing view that 5hmC predominantly marks accessible chromatin, MNase fractionation combined with 5hmC capture shows that the vast majority of 5hmC resides in MNase-protected DNA, consistent with nucleosome association. Nonetheless, we found that small subsets of 5hmC were more MNase-sensitive (accessible 5hmC) or -resistant (protected 5hmC) relative to the common pool. With chromatin-state annotations, we further separated these subsets where the accessible 5hmC aligned with promoter/enhancer states and the protected 5hmC aligned with mixed permissive-repressive marks. Together, these results indicate that 5hmC is predominantly found in the protected fraction and that 5hmC is heterogeneous, with context-dependent relationships to chromatin and transcription. These MNase-biased subsets may also represent transitional chromatin states, a possibility that remains to be tested.

Motif analysis suggests that the two differential 5hmC subsets reside in distinct TF environments. Accessible 5hmC is enriched for PU.1 and IRF motifs, consistent with prior studies by us and others showing that PU.1, which can partner with IRF, recruits TET enzymes to B lineage enhancers to promote local DNA demethylation and accessibility^8,28^. In contrast, protected 5hmC is enriched for the motifs of several KLFs, including KLF2 and KLF4, which are essential for enforcing quiescence and trafficking programs in naïve B cells and can bind to methylated CpG and nucleosomal DNA^29,30^. Notably, KLF4, the most well-characterized KLF family member, has been shown to recruit TET to promote DNA demethylation on nucleosomes^19^. Consistent with the fact that the protected 5hmC is enriched in bivalent-like chromatin state that is potentially related to cell differentiation, KLF5 has been shown to regulate bi-potential cell fate differentiation in embryonic stem cells^31^. These observations support a model in which TFs like PU.1 and IRF help establish the accessible-5hmC pool in B cells. In contrast, KLF factors could promote TET activity on nucleosome-bound DNA via their pioneer-factor-like activity and may promote the bivalent-like state^19^. Note that the bivalency-like state here differs from the classical bivalent chromatin that is typically marked by H3K4me3 and H3K27me3. How the H3K9 bivalent-like state enriched in the protected 5hmC is regulated, and how its role in gene regulation remains unclear.

Across tissues and systems, gene-body 5hmC shows a robust positive relationship with gene expression levels that is superior to the gene-body 5mC, which also tracks with gene expression levels^32,33^. Recent studies using deep learning showed that 5hmC alone can classify gene expression states^24,34^. Consistent with Gonzales-Avalos *et al.*, we found that the promoter-proximal 5hmC has better correlation with gene expression^24^. Our finding that promoter-proximal, MNase-sensitive (accessible) 5hmC correlates more strongly with transcription than unstratified 5hmC may be able to enhance the performance of these learning models.

Several limitations exist for our current study. For instance, to completely remove accessible DNA, we used a higher MNase concentration to digest the chromatin to obtain mononucleosomes. The 5hmC associated with MNase-sensitive fragile nucleosomes was likely excluded from our nucleosomal DNA fractions and instead represented as accessible 5hmC^35^. The degree to which 5hmC is associated with these fragile nucleosomes remains to be determined. Besides the MNase-based method, future work may further distinguish different particle types of nucleosomes using other methods such as NOMe-seq (Nucleosome Occupancy and Methylome Sequencing). While these considerations may temper mechanistic interpretation, they do not alter our central finding that 5hmC predominantly resides in the MNase-protected fraction and that the accessible and protected subsets align with distinct chromatin states.

In summary, our data suggest that 5hmC is predominantly associated with MNase-resistant nucleosomes in naïve B lymphocytes. We identified two functionally distinct subsets of 5hmC according to their relative sensitivity to MNase, and these subsets differ not only by their associated chromatin states but also by their relationship with gene expression and potential regulatory networks.

## Materials and Methods

### Animals

All animal experiments were approved by the IACUC committee at the Ohio State University (protocol number 2020A00000055-R1). C57BL/6J mice of both sexes (6-12 weeks; Jax#000664) were purchased from Jackson Laboratory (Bar Harbor, ME).

### Isolation of primary mouse B cells

Mouse B cells were isolated as we previously described^36^. Spleens from B6 mice were isolated and pressed against a strainer to obtain total splenocytes. Erythrocytes were lysed using ACK lysing buffer (Gibco). B cells were isolated from 50 million splenocytes with EasySep Mouse B Cell Isolation Kit following the manufacturer’s instructions (Stemcell Technologies). Cell purity was confirmed using flow cytometry and was >95%.

### MNase Digestion

Naive B cells were pelleted by centrifugation at 400 *xg*, 4LC for 5 minutes and then resuspended to 20 x10^6^ cells/mL in 1x swelling buffer (10 mM Tris pH 7.5, 2 mM MgCl_2_, 3 mM CaCl_2_). After 5 minutes of incubation on ice, cells were spun at 400 *xg* for 10 minutes. Supernatant was discarded, and the cell pellet was resuspended in the same volume of 1x swelling buffer with glycerol (1x swelling buffer, 10% glycerol). Another volume of lysis buffer (1x swelling buffer, 10% glycerol, 1% NP40) was slowly added to the sample with agitation. After 5 minutes of incubation on ice, the cells were pelleted by centrifugation at 600 *xg* for 5 minutes. The supernatant containing the cytoplasmic fraction was discarded. A gradient of micrococcal nuclease (MNase; NEB) was arranged by a 3x serial dilution from 50x to 1350x for each reaction with 1x10^6^ cells in swelling buffer (10 mM Tris pH 7.5, 2 mM MgCl_2,_ 3 mM CaCl_2_). Cell nuclei and prepared MNase reaction mixes were heated to 37LC for 2 minutes before mixing. The reaction mixture was incubated at 37LC for 10 minutes before prompt quenching with a 10x Stop Buffer (200 mM EDTA pH 8.0, 40 mM EGTA pH 8.0) to neutralize the MNase reaction. DNA was purified using the Zymo DNA Clean and Concetrator-5 (Zymo Research) and quantified using Qubit dsDNA HS assay (ThermoFisher). Digestion efficiency was verified using the 2200 Tapestation and D1000 ScreenTape kit (Aglient Technologies). For large-scale complete nucleosome digestion, we used 50x diluted MNase.

### DNA sonication

Genomic DNA (0.5 to 1μg) in 100 μL 1xTE was sonicated to 150-200 bp using a Bioruptor Pico 2 (Diagenode; 30 cycles of 30s on, 30s off in 4LC circulating water). Sonicated DNA was purified using 1.6x Ampure beads and quantified using Qubit DNA HS assay (ThermoFisher).

### 5hmC enrichment sequencing (hMe-Seal)

5hmC in sonicated and nucleosomal DNA was analyzed as we previously described^36^. Briefly, DNA was end-repaired using Kapa HyperPrep (Kapa Biosystems), followed by adapter ligation. Unmodified complementary DNA strands were synthesized using primer extension with Klenow Exo^−^ and a primer specific for the 5’ of the ligated adapter. 5hmC was glycosylated with T4 beta-glucosyltransferase (T4-BGT; ThermoFisher) and 3 nmol of UDP-N_3_-glucose (Jena Bioscience) at 37LC for 1 hour. DBCO-PEG4-Biotin (20 nmol) was added to the reactions for two additional hours at 37LC where DBCO reacts with the azide group via a CLICK chemical reaction mechanism. DNA was purified with the DNA Clean and Concentrator 5 kit (Zymo), and the biotinylated DNA fragments were enriched using T1 Streptavidin Dynabeads (ThermoFisher) on a thermoshaker for 30 minutes (1300 rpm, 25LC). Beads were collected using a magnet and washed twice with 1x BW buffer (1M NaCl, 10 mM Tris-HCl pH 7.5, 1 mM EDTA pH 8.0, 0.02% Tween-20) on a thermoshaker for 5 minutes (1500 rpm, 25LC). Beads were washed twice with TE20 (10 mM Tris-HCl pH 8.0, 1 mM EDTA pH 8.0, 0.02% Tween-20) on a thermoshaker for 5 minutes (1500 rpm, 25LC). The biotinylated DNA on the beads was denatured for 10 minutes with 0.1 N NaOH to release the synthesized complementary strand, followed by neutralization using 1 M acetic acid. The released DNA strands in the supernatant were collected and used for library generation.

### Library preparation and Illumina sequencing

Sequencing libraries were generated using the NEBNext Ultra™ II DNA Library Prep Kit for Illumina (NEB) as we previously described^36^. Libraries were barcoded using NEBNext Multiplex Oligos for Illumina (96 Unique Dual Index Primer Pairs; NEB), analyzed using DNA 1000/5000 tapes with TapeStation (Agilent), and quantified with Qubit dsDNA HS Assay Kit (ThermoFisher). Libraries were pooled and sequenced using Illumina Nova-seq 6000 at the Institute for Genomic Medicine at the Nationwide Children’s Hospital (Columbus, OH).

### HPLC-MS 5mC and 5hmC Quantification

Subnucleosomal and sonicated genomic DNA were prepared as described above. For quantification of global cytosine modifications, 1 µg of DNA was enzymatically hydrolyzed using DNA Degradase Plus (Zymo Research) in a 25 µL reaction at 37 °C for 8 hours, followed by enzyme inactivation at 70 °C for 30 minutes. To generate a standard curve for absolute quantification, known concentrations of 2′-deoxycytidine, 5-methyl-2′-deoxycytidine, and 5-(hydroxymethyl)-2′-deoxycytidine (all from Cayman Chemical) were prepared at 5, 25, 125, 250, 500, and 1,000 ng/mL. Chromatographic separation was performed using an Atlantis T3 column (100 Å, 3 µm, 2.1 mm × 100 mm; Waters) on a ThermoFisher Scientific UltiMate™ 3000 HPLC system. The mobile phase consisted of phase A: water with 0.1% formic acid (ThermoFisher), and phase B: acetonitrile with 0.1% formic acid (ThermoFisher), delivered at a flow rate of 0.2 mL/min. A 5 µL volume of each sample was injected. The gradient elution was programmed as follows: 0% B from 0–1 minute; linear increase to 100% B from 1–5 minute; hold at 100% B from 5–7 minute; decrease to 0% B at 9 minute; re-equilibrate at 0% B from 9–10 minute. The column was maintained at 40 °C throughout the run. Mass spectra were acquired on an Orbitrap Exploris 480 mass spectrometer (ThermoFisher) operating in positive ion mode. Raw spectra were processed using the ThermoFisher Xcalibur software. Retention times and m/z values were verified manually in negative mode, and peak areas were initially integrated using the Quan Browser’s processing method, followed by manual adjustment for both standards and samples. Peak area data were exported and used to generate linear standard curves for 5hmC, 5mC, and C, which were then used to interpolate concentrations in the biological samples.

### Bioinformatics Analyses

#### Data Sources

Deep sequencing datasets used for correlative bioinformatic analysis were sourced from accessions SRA: SRP029721^37^ and GEO: GSE116208^7^, GSE82144^26^.

#### ChIP and 5hmC analysis

Paired-end reads were quality checked (FastQC v0.11.8)^38^ and trimmed (cutadapt v 4.1)^39^ before mapping to the mouse genome (mm10) using Bowtie2 (2.4.2)^40^. Unmapped reads and duplicates were removed using Samtools (v1.10)^41^. BigWig files were generated using bamCoverage (--normalizeUsing RPKM) using deepTools (v3.5.1)^42^. Averaged bigwigs were generated using wiggletools (https://github.com/Ensembl/WiggleTools). Peaks per biological replicate were called with MACS2^43^ (-g mm -q 0.05 –keep-dup all). Differential analysis for 5hmC data was conducted with DiffBind using default DESeq2 settings and peak files generated by MACS2^44^. Initial visualization of peaks on scatterplots used both an adjusted p-value of 0.05 and an absolute fold change of 1.5 or greater for differential regions. Downstream analyses afterwards used an adjusted p-value of 0.05 and an absolute fold change greater than 0 to maximize protected and accessible 5hmC regions compared. For comparison, random regions were generated without overlap (Bedtools 2.30.0)^45^. Heatmaps were made using deepTools (computeMatrix reference-point – referencePoint center -bs 100 -a 3000 -b 3000 -sortRegions keep).

#### T4gt Spike-In Comparisons

Paired-end reads not aligned to mm10 were trimmed, mapped, and filtered as mentioned above against the T4 bacteriophage genome (NC_000866.4). The mapped T4 read counts were normalized using total mapped reads of 5hmC enrichment to mm10.

#### DNA methylation analysis

Single-end reads were mapped to bisulfite-converted mouse genomes (mm10) using Bismark and Bowtie 2 (0.22.1)^46^. Aligned reads were deduplicated, and methylation levels were analyzed using Bismark (deduplicate_bismark and bismark_methylation_extractor, respectively). The resulting bedgraph of methylation rates per CpG was processed into a symmetrized format using DNMTools^47^ before a 10x CpG coverage filter was applied. The formatted bedgraph was converted into a bigwig using UCSC kentUtils *bedGraphToBigWig*. Profile plots of methylation over the indicated regions in the figures were made using deepTools.

#### Chromatin accessibility analysis

Paired-end reads were processed similarly to ChIP and hmC reads. After deduplication, reads were shifted to account for Tn5 insertions and reformatted to the minimal BEDPE format. Peaks were called with MACS2 as broadpeaks (-g mm -q 0.05 –keep-dup all –broad). BigWigs and heatmaps were generated as above with deepTools.

#### RNA-seq integration

The analyzed RNA sequencing data in B cells were obtained from our previous study^7^. The transcripts per million (TPM) values per gene were averaged over replicates. To assign expression quartiles, we obtained the expression levels of all genes. Genes above the threshold were assigned quartiles (lowest 25% in Q1, highest 25% in Q4) in relation to the TPM of all other remaining genes. Genes with TPM of 0 or less were considered to be not expressed (NE). Random regions and differential 5hmC peaks were annotated using HOMER (v5.1)^48^, and the expression of the nearest genes was tabulated.

#### Motif analysis

Transcription motifs enriched at differential 5hmC regions were determined using HOMER with *findMotifsGenome.pl* (v5.1), with the random regions and common 5hmC regions as the background^48^. The top 5 significant motifs were shown.

## Acknowledgements

We want to thank Drs. Partick Collins, Eugene Oltz, Mark Parthun, and Edahi Gonzales-Avalos for advice and comments; members of the Lio and Ma labs for technical support and suggestions. Drs. Gong Wu and Jiangjiang Zhu for their assistance with HPLC-MS.

This research is supported by the Department of Microbial Infection and Immunity, the Pelotonia Institute for Immuno-Oncology at OSU, NIH NIGMS R35 (R35GM151110), and Gabrielle’s Angel Foundation (to J.L.). H.C. was supported by the Pelotonia postdoc fellowship and by the American Heart Association postdoctoral fellowship.

---

**Supplementary Figure 1.**
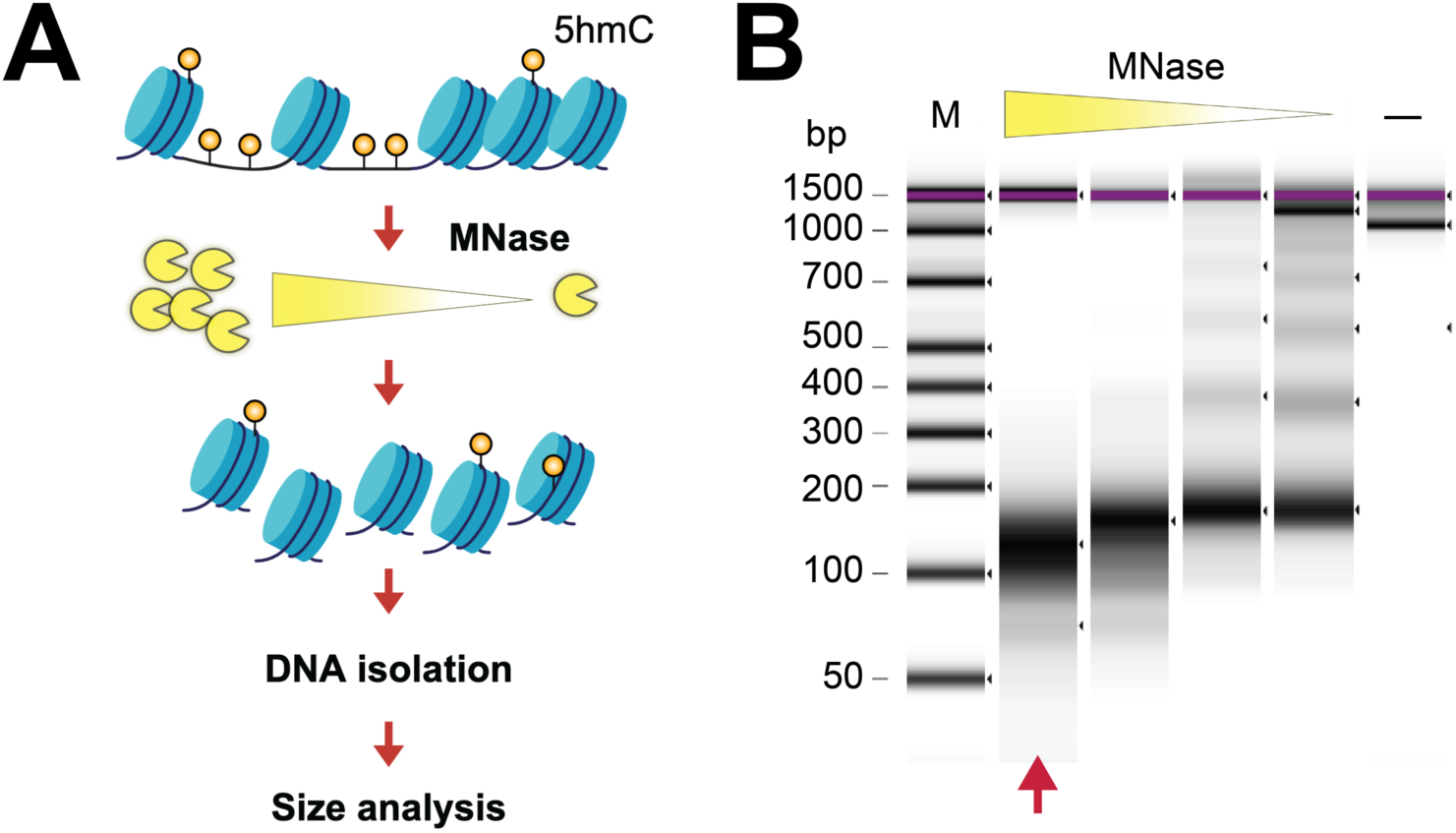
Titration of MNase digestion. **(A)** Schematic for the MNase titration experiment. Nuclei were digested with titrated levels of MNase. DNA was purified, and the fragment size was analyzed using TapeStation. **(B)** Electrophoresis diagram of MNase titration. From left to right, the MNase dilutions are 50x, 150x, 450x, and 1350x. DNA isolated from undigested nuclei was shown as a control (**-**). M, DNA markers. The red arrow indicated the chosen condition for subsequent experiments. Representative data from at least three experiments are shown.

**Supplementary Figure 2.**
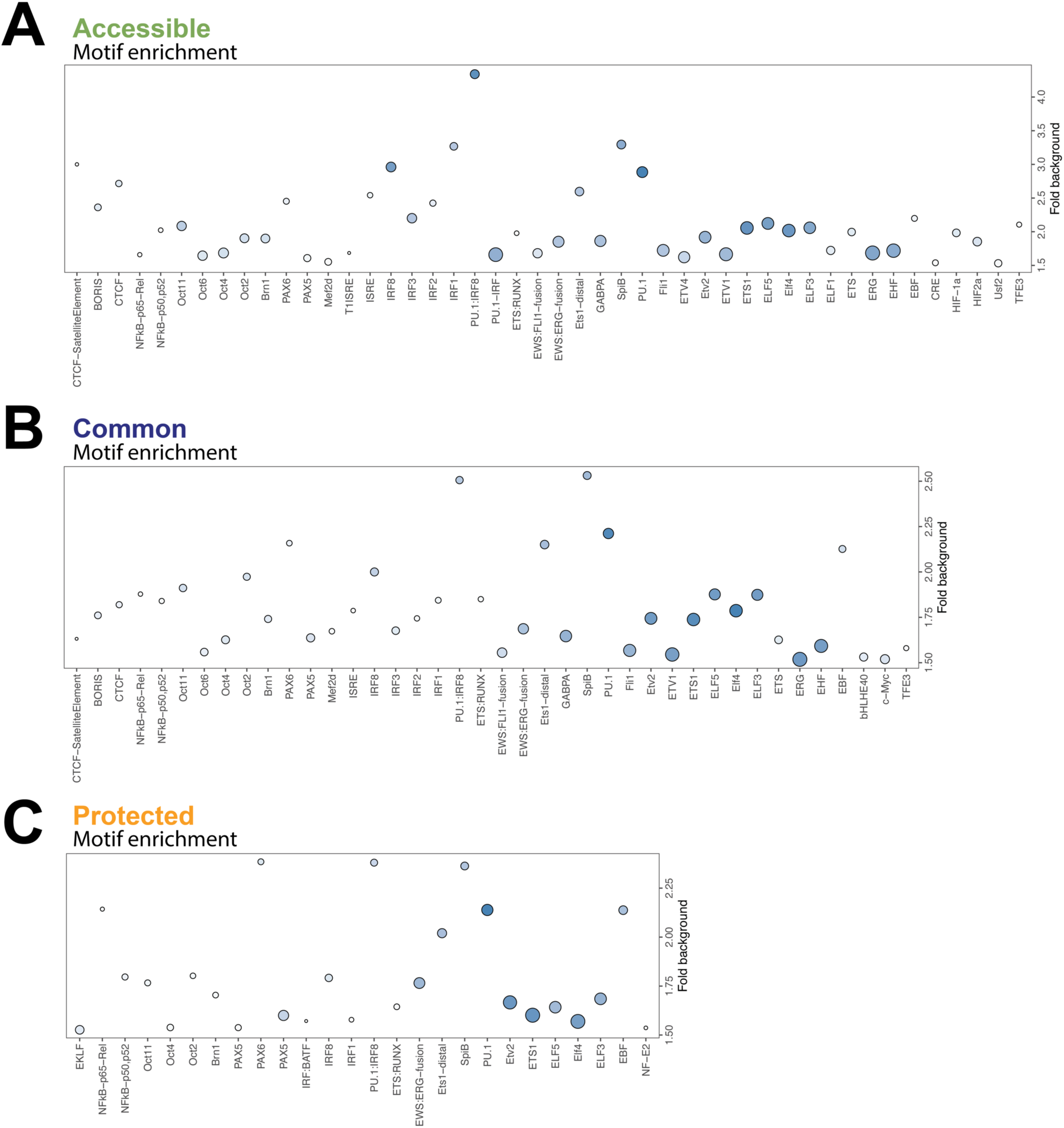
Motif enrichments at 5hmC regions. Motifs enriched at **(A)** accessible, **(B)** common, and **(C)** protected 5hmC regions were analyzed using random genomic regions as the background. The enrichment levels were shown.

**Supplementary Figure 3.**
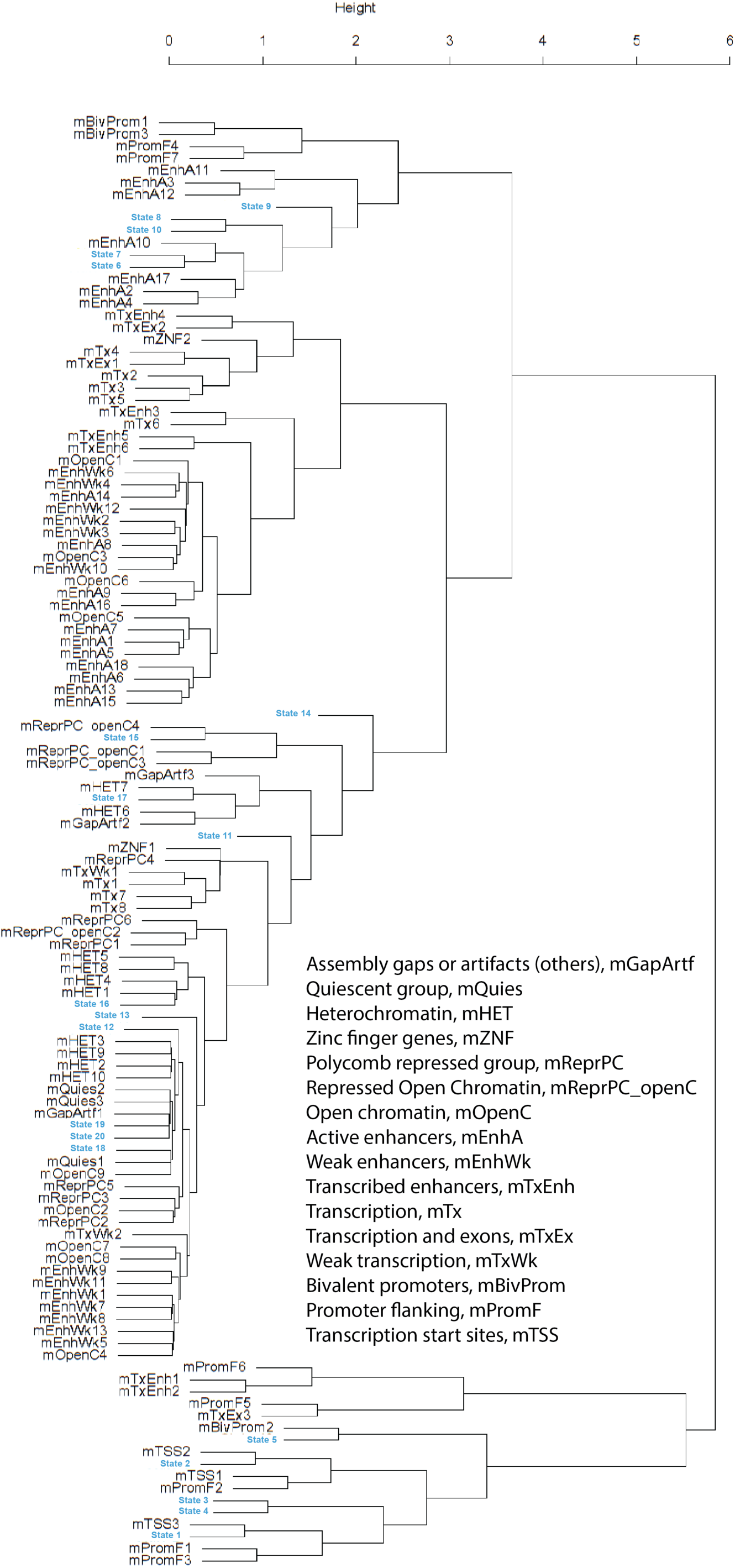
Informed annotation of chromatin states. Hierarchical clustering tree of generic 20 ChromHMM states against 100 universally labeled mm10 chromatin states. Using the emission probabilities of ChIP-seq data common between the generated 20 ChromHMM states and the 100 universal mm10 chromatin states, hierarchical clustering was used to identify the most probable label for each of the generated states by Manhattan distance.

**Supplementary Table 1. Deep sequencing data sources.**

**Supplementary Table 2. ChromHMM training datasets.**

